# Metabolons, quinary structure, and domain motion: enzyme choreography in the cytoplasm

**DOI:** 10.1101/2022.09.13.507800

**Authors:** Premila P. Samuel Russell, Meredith M. Rickard, Taras V. Pogorelov, Martin Gruebele

## Abstract

How do enzymes form metabolons inside cells? To answer that question, we created an all-atom model of a section of the human cytoplasm and simulated it for over 30 microseconds. Among other proteins, nucleic acids, and metabolites, the model contains three successive members of the glycolytic cycle: glyceraldehyde-3-phosphate dehydrogenase (GAPDH), phosphoglycerate kinase (PGK), and phosphoglycerate mutase (PGM). These enzymes interact to form transient, but long-lived, multi-enzyme complexes with characteristic lifetimes in the 1 to 5 μs range, thus modeling the functional metabolon structures that facilitate compartmentalization of metabolic pathways and substrate channeling in cell. We analyze the quinary structure between enzymes down to the formation of specific hydrogen-bonded interactions between side chains, together with the movement, in concert, of water molecules in or out between interacting amino acids to mediate contact formation and dissolution. We also observed large-scale enzymatic domain motion that has been proposed to convert between substrate-accessible and catalytically functional states: a direct hinge-bending motion of up to 28° changes the relative orientation of the N- and C-terminal domains of PGK, causing the initially open, and presumably inactive, conformation of PGK to sample both “semi-closed” and “closed” conformations. Although classical molecular dynamics (MD) cannot simulate enzymatic activity, closed structures are the functionally active forms of PGK, and their equilibrium with open structures opens the door for future quantum mechanics/molecular mechanics (QM/MM) and other reactive simulations of the cytoplasm.

## Introduction

The cellular environment influences enzyme function in multiple ways, from compartmentalization to substrate diffusion. In particular, protein-protein interactions can be viewed as having a crowding (repulsive)^1^ and sticking (attractive)^2^ components.

Crowding and sticking can lead to the emergence of large-scale domain motions and structural rearrangements in an individual enzyme,^3^ as well as the emergence of transient functional interactions, called quinary structure^4^, to favor substrate channeling within the multi-enzyme complexes known as metabolons^5,6^. Crowded *in vitro* model environments have been seen to favor enzymatically active compact structures of phosphoglycerate kinase (PGK) and adenylate kinase^3,7^. Canonical examples of enzyme-enzyme quinary interactions have been seen along the glycolysis pathway, such as between GAPDH and PGK, which have been shown to associate weakly in the cytoplasm ^8,9^ with an effective dissociation constant *K*_*d*_ of 14±3 μM, presumably improving substrate processing by reducing the average distance for substrate diffusion.^10^ Finally, experiments have shown that protein diffusion decreases in cells, indicating clustering when the interaction of a protein with its environment is increased^11–13^. The weakness of quinary structure between enzymes in metabolons is critical, because it further allows enzymes to dissociate quickly and associate with other binding partners for multiple functions, including with neighboring enzymes on the metabolons^14^ or having enzymes act in signaling chains.^14^ The GAPDH/PGK pair, in particular, has been seen organized into such higher order multi-enzyme structures along the glycolytic pathway^15,16^.

In-cell molecular dynamics simulations^17^ have recently reached the multi-μs time scale necessary to observe multi-protein clustering^18,19^ and transient protein-protein interactions in the bacterial cytoplasm^20,21^. Here, we build an all-atom model of a segment of the human cytoplasm, and over the course of 31.2 μs of MD simulation, we observe the transient assembly, disassembly, and re-assembly of pairs of enzymes into a tri-enzyme quinary structure, where the pairs are either GAPDH/PGK, GAPHD/PGM, or PGK/PGM.

The quinary structure is analyzed from general pairwise interactions of protein surfaces all the way down to individual water molecules that mediate the formation or dissolution of protein surface interaction. We are also able to resolve how crowding directs large-scale hinge-bending motion of the enzyme PGK, supporting the idea based on *in vitro* studies^7^ that cytoplasmic crowding can push enzymes into catalytically active conformations.

## Results and Discussion

### A full-atom model of a slice of the U-2 OS cytoplasm

Our initial modeling studies on protein-protein sticking were done in a bacterial cell cytoplasm environment, mimicking *E. coli*. However, bacterial and mammalian cellular conditions differ significantly. For example, metabolite concentrations of glutamate are three times higher in *E. coli* than in *H. sapiens*. Conditions in the cytoplasm can also vary greatly across human cells with cell type, developmental stage, and circadian rhythm ^22–26^.

In order to investigate mammalian cellular sticking and crowding mechanisms leading to metabolon formation, we constructed a small human cytoplasm model to closely mimic proteomic and metabolomic data of the U-2 OS osteosarcoma cell line^10^. In our cytoplasm model, with one notable exception (GAPDH)^9^, we opted to mostly include proteins that were relatively small (≤ 50 kDa) to ensure free translation of the proteins throughout the model. We also included three copies of the very small bacterial protein B (PrB), a model fast-folding protein. The resulting computationally relatively inexpensive model size of 203,903 atoms enabled us to carry out 31.2 μs of simulations, sufficient to sample a hierarchy of cellular-level dynamics.

Both GAPDH and PGM in our model are of human origin, while *saccharomyces cerevisiae* yeast PGK3 R65Q/F333W (65 % sequence identity with human PGK1) was used to allow direct comparison with previous simulations and in-cell experiments of GAPDH-PGK interactions and PGK domain rearrangement^7,9,27–30^. Anaerobic glycolytic pathways are conserved between human and yeast cell lines^31^. Due to high sequence identity, we still were able to investigate several pairs of wild-type co-evolving surface residues in GAPDH/PGK and PGM/PGK pairs for the functional formation of quinary interactions^32^.

We used two different experimental studies to determine a consensus metabolite abundance table for human cells^23,24^. In the cytoplasm model, 44 different species of metabolites were sampled by Monte Carlo selection from the abundance table, ranging from the most abundant (glutamate, 32 mM) to 77^th^ most abundant (GMP, 0.1 mM). A comparison of the metabolite concentrations represented in the *H. sapiens* model and the consensus experimental concentrations is included in supplement (SI Table 1).

The cellular composition of the cytoplasm model is ∼280 mg/mL of macromolecules, ∼175 mM metabolites, and ∼190 mM inorganic ions. These concentrations are lower than for our previous *E. coli* model due to lower experimentally measured metabolite and inorganic concentrations determined for human U2-OS cells^25^ and similar human cell lines.

### Short-lived universal and long-lived specific sticking interactions

We computed the temporal statistics of pairwise protein sticking in the cytoplasmic model (excluding PrB), in analogy to our previous study of the bacterial cytoplasm, which did not contain quinary pairs. Fig. 2 shows the probability distribution *P* of contact times *Δt* for a minimum surface area of 5 nm^2^ on a ln(*P*) vs. ln(*Δt*) scale. Below about 10 μs, we observe the same slowly decreasing contact probability with contact time that was fitted to a *t* ^*-1*^ power law in our bacterial study. There, the scaling behavior was assigned to a previously proposed universal “birth-death” mechanism^33^ that also applies to protein sticking interactions. At *Δt* > 10 μs, where no significant contacts were observed previously, the distribution rebounds in our human cell model. This is mainly due to long-lived interactions between the three glycolytic enzymes, as well as these enzymes with other cytoplasmic proteins such as cyclophilin and HAH, a protein extensively studied by in-cell NMR to reveal its sticking dynamics^13,34^.

### Finding order in chaos: metabolon complexes among the sticking interactions

Protein-protein contact events for all protein pairs in the cytoplasm model were measured in terms of the solvent accessible surface area (SASA) decrease with contact site formation, as previously done for our bacterial cytoplasm models. The results in Fig. 3 show a very wide dynamic range from long-lasting contacts to highly transient interactions^20^. While our model underwent significant protein diffusion (Fig. 1), of course a lack of interaction cannot be interpreted as that protein pair not interacting *in vivo* because the 31.2 μs of sampling are still far from ergodic regarding the mixing of protein locations.

**Fig. 1.**
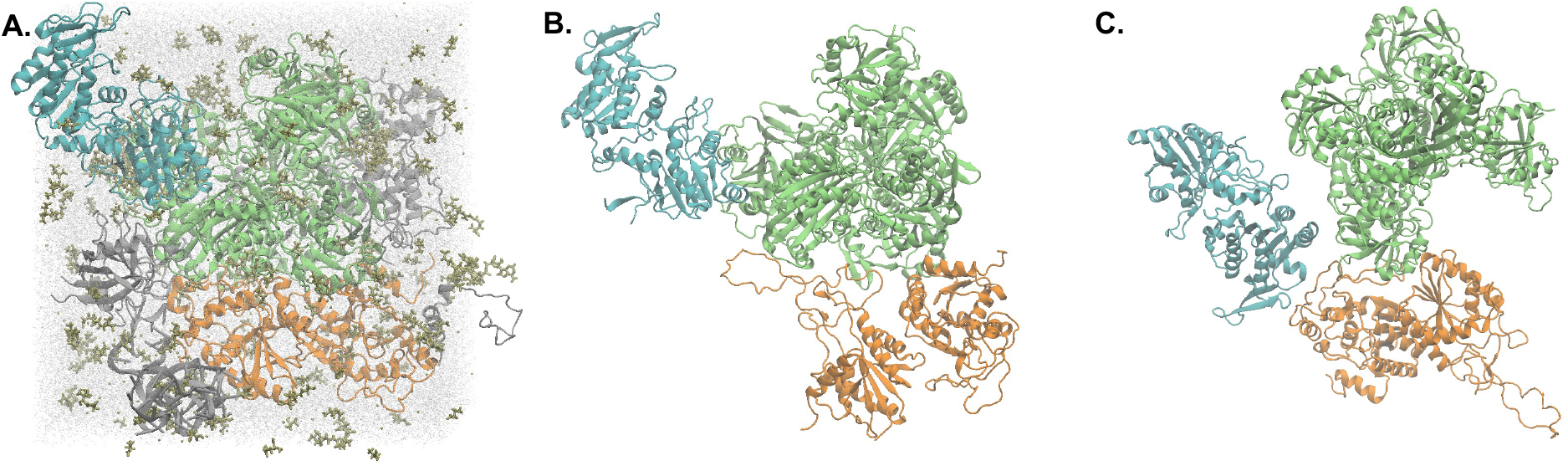
Glycolytic enzymes in a mammalian cell cytoplasm model. (A) Snapshot of the U-2 OS cell cytoplasm atomic model at time 0 μs (after equilibration is completed) of the MD production run. Macromolecular backbones are represented as cartoons, with color depictions of yeast PGK as cyan, human GAPDH as lime, human PGM as orange, and all other macromolecules as silver. Water molecules are represented as transparent silver-colored CPK models, and ligand and ions are represented as tan-colored licorices. (B) Snapshots of PGK, GAPDH, and PGM forming transient glycolytic complexes at 22 μs, and (C) at 23.9 μs along the MD trajectory, showing significant diffusion and changes in contact formation.

We observed among these protein sticking events the three glycolytic enzymes in the cytoplasm organizing together to form evolving conformations of multi-enzyme complexes (Fig. 1B and 1C) lasting up to several microseconds. Fig. 3 shows such interactions forming, dissolving, and, in some cases, re-forming between GAPDH-PGM, PGK-GAPDH, and to a lesser extent PGM-PGK. As a structural highlight, in Fig. 1B at 22 μs we observe GAPDH interacting simultaneously with PGK and PGM. About 2 μs later, at 23.9 μs, the interaction between PGK and GAPDH has been broken, while a new interaction has formed between PGK and PGM. The high *K*_*d*_ for PGK:GAPDH disassociation (∼14 μM)^8,9^ enables PGK to break contact with GAPDH and form contacts with PGM. Several past studies have identified different combinations involving these three enzymes and their homologs in complex formations^15,16^, including GAPDH also short-circuiting PGK to provide substrate to a PGM homolog.

### Quinary interaction between glycolytic enzymes at the single residue level

Transient association and dissociation events at the residue level (with a distance cutoff of 0.45 nm) were analyzed between all three proteins and summarized in Figs. 4 and 5.

We begin with PGK-GAPDH. Experiments have shown that PGK and GAPDH can form cross-species quinary structure^8–10^ between yeast PGK and human GAPDH with a measured dissociation constant of 14 ± 3 μM, which is on the order of PGK and GAPDH concentrations in U2-OS cells^9^. Here we see that the PGK N-terminal domain forms the largest number of transient contacts with GAPDH subunit 2 (GAPDH2) (Fig. 4A). Of the 43 longest-lived residue-residue interactions (occurring for at least 10% of the of the simulation), all but 5 residue-residue pairs involved at least one residue conserved in both yeast and human species (Fig. 4E). Some residue pairs involve more than one pair of interacting atoms at a time, for example between the ammonium group on Lys145 of GAPDH2 and the carboxyl group of Asp143 on PGK (see discussion of H-bond network below).

For each specific inter-chain contact, a larger surrounding protein patch emerged as a quinary structure hallmark. For example, for the PGK/GAPDH2 interaction, PGK residues numbered 96-108 and 130-143 formed two such patches. Many of the inter-residue interactions between these patches lasted for over 5 μs (Fig. 4A,B), and we take this as a characteristic lifetime for the quinary structure observed here. These patches are responsible for the increased probability of the longest protein-protein contact time intervals in Fig. 2.

**Fig. 2.**
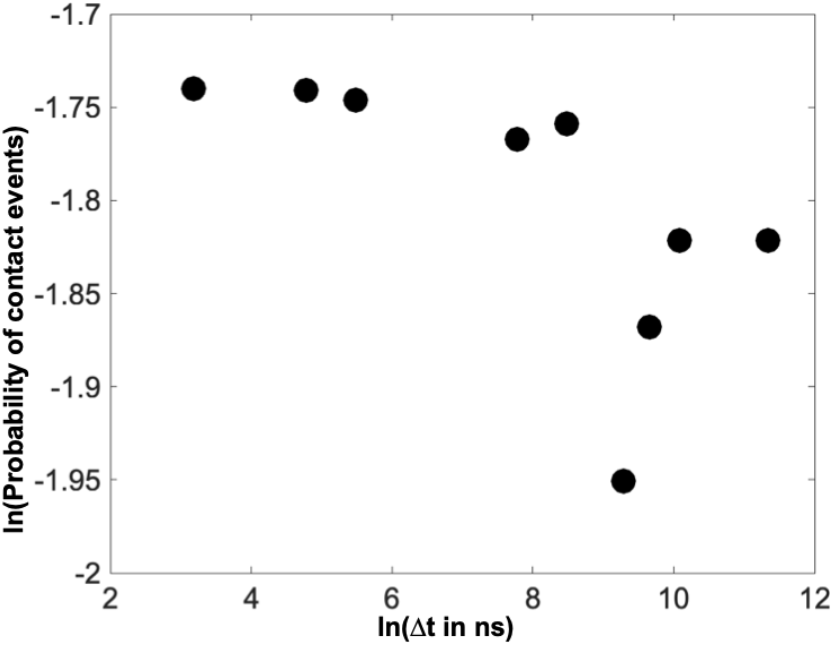
Contact time probability distribution for protein pairs in the model U-2 OS cytoplasm. Only interactions with a common surface area ≥ 5 nm^2^ were included in the distribution, similar to ref. ^20^. The probability initially decreases, as observed for the bacterial cytoplasm as well, but the human cell simulation with its coupled glycolytic enzymes shows a strong rebound above 10 microseconds. Homo-oligomers (e.g. the GAPDH tetramer) are not included in the statistics.

PGK’s N-terminal domain holds the binding site for 1,3-bisphosphoglycerate, while the C-terminal domain binds ADP^14^. Published machine learning-based predictions of the PGK/GAPDH interface have shown that both N- and C-terminal PGK domains interact with GAPDH, supporting a bi-domain mechanism of interaction^35^. While interactions between GAPDH subunit 3 (GAPDH3) and yeast PGK’s C-terminal domain only involved two residue-residue interactions over 5 μs (Fig. 3B), they were indeed concurrent with some of the interactions between GAPDH2 and the PGK N-terminus. It has been hypothesized that a metabolon with more than one GAPDH per PGK evolved in order to channel 1,3-bisphosphoglycerate substrate efficiently to PGK enzymes and thus reduce time-consuming metabolite diffusion between steps in the reaction pathway^36^.

**Fig. 3.**
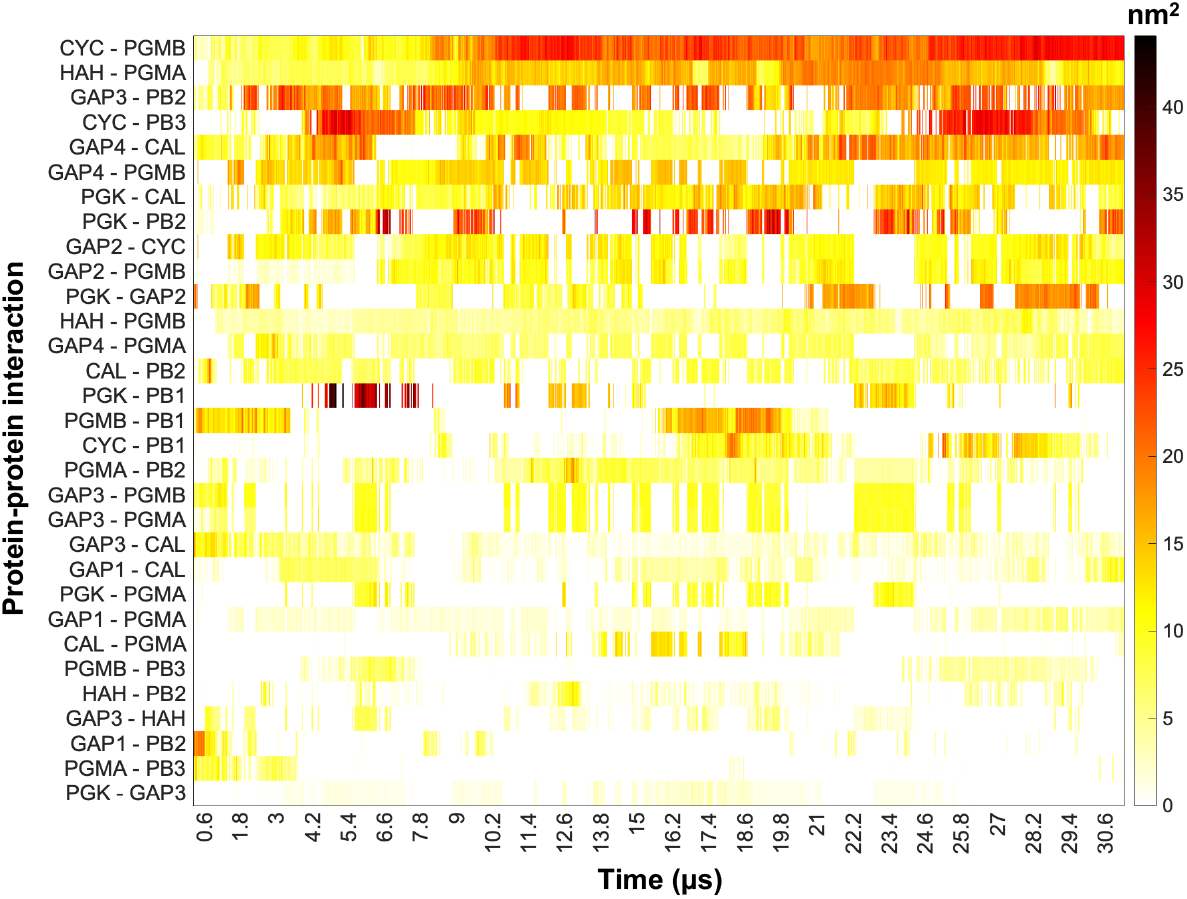
Heat map of the contact surface area for the largest protein-protein sticking interactions in the mammalian cell. Interaction between subunits or copies of a same type of protein are not shown here. Protein abbreviations are as following, with one subunit/copy example given per multiple copies of a protein chain: CYC, cyclophilin G; CAL, calmodulin; HAH, copper transport protein HAH1; GAP1, 1^st^ subunit of GAPDH; PGMA, subunit A of PGM; and PB1, protein B copy 1. Each interacting protein pair shown here have total interacting area of ≥ 87.9 nm^2^ across the simulation trajectory.

We observe similar interactions between PGK and PGM (Fig. 4C,F). Although not studied in detail experimentally yet, such transient structure could likewise facilitate the passing of 3-phosphoglycerate between these two enzymes.

**Fig. 4.**
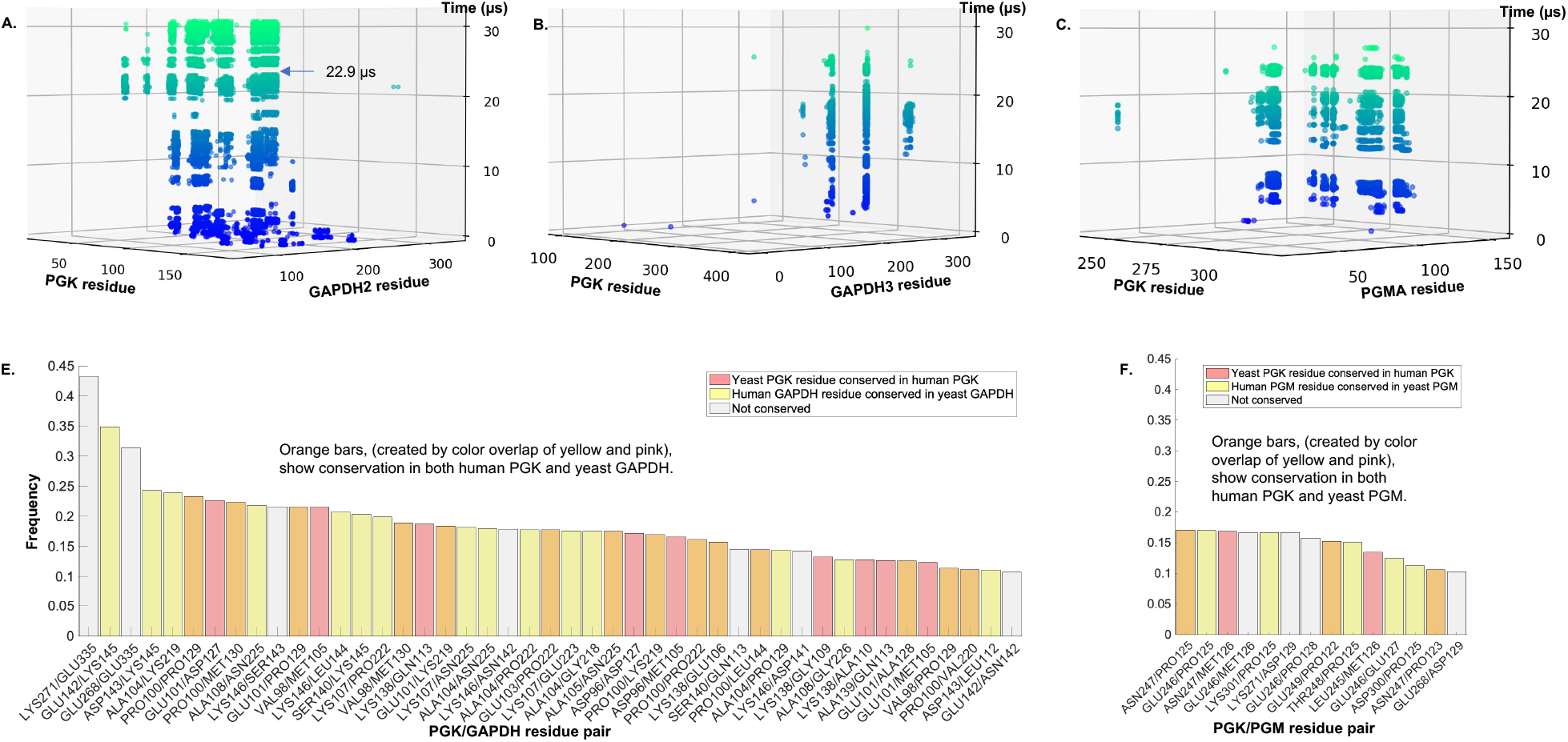
Residue-specific contacts formed between yeast PGK and human glycolytic enzymes during the mammalian cytoplasm simulation. Time traces for residue-residue contacts formed between yeast PGK and (A) GAPDH subunit 2; (B) GAPDH subunit 3; and (C) PGM subunit A. Each contact is shown as a circle, undergoing color change from blue to light green with increasing time. Residue-residue contact frequency bar charts for most frequent (≥ 10%) quinary interactions between (E) PGK and GAPDH (either 2^nd^ or 3^rd^ subunit) pairs; and (F) PGK and PGM (subunit A) pairs. The distance cutoff for the residue-residue contacts in (A)-(F) was 0.45 nm. The bar plots are further colored to reflect cross-species (human/yeast) enzyme residue conservation of homologs: pink, PGK residue is conserved; yellow, either GAPDH (E) or PGM (F) is conserved; orange, interacting residue pairs are conserved across human and yeast; and white, no cross-species residue conservation. The yeast PGK3 R65Q/F333W variant used in our simulation was aligned to human PGK 1 with 65.22% sequence identity; human GAPDH was aligned simultaneously to the 3 variants of GAPDH in yeast: GAPDH 1 with 64.44% sequence identity, GAPDH 2 with 64.74% sequence identity, and GAPDH 3 with 65.05% sequence identity; and human PGM was aligned to yeast PGM 1 with 51% sequence identity. A residue is marked here as conserved if the corresponding residue occurring cross-species either is same or similar. We grouped similar amino acids as following: GAVLI, FYW, CM, ST, KRH, DE, and NQ. For GAPDH alignment, if the human GAPDH residue was conserved in at least 2 yeast GAPDH variants, we marked the residue here as conserved. All interacting enzyme subunit pairs were included here in analyze only if they had total interacting area of ≥ 87.9 nm^2^ across the simulation trajectory.

The largest number of interactions across the three enzyme pairs was observed for PGM:GAPDH, and is summarized in Fig. 5. These two enzymes are not sequential along the “normal” glycolytic pathway. Interestingly, however, a homolog of PGM (with ∼53% sequence identity) in human erythrocytes called bis-PGM is involved in a special glycolytic step involving the direct passing of substrate between GAPDH and bis-PGM. This step is called the ‘Rapoport-Luebering Shunt’ and is unique to erythrocytes^37^. Fig. 5D highlights residue pairs contributing to the longest-lived contacts between GAPDH and PGM for which the interacting PGM residues are also conserved in erythrocyte bis-PGM. Additional contacts with interaction frequency between 30% to 10%, and the non-conserved contacts, many of which are equally long-lived, are shown in supplementary information Fig. S1.

**Fig. 5.**
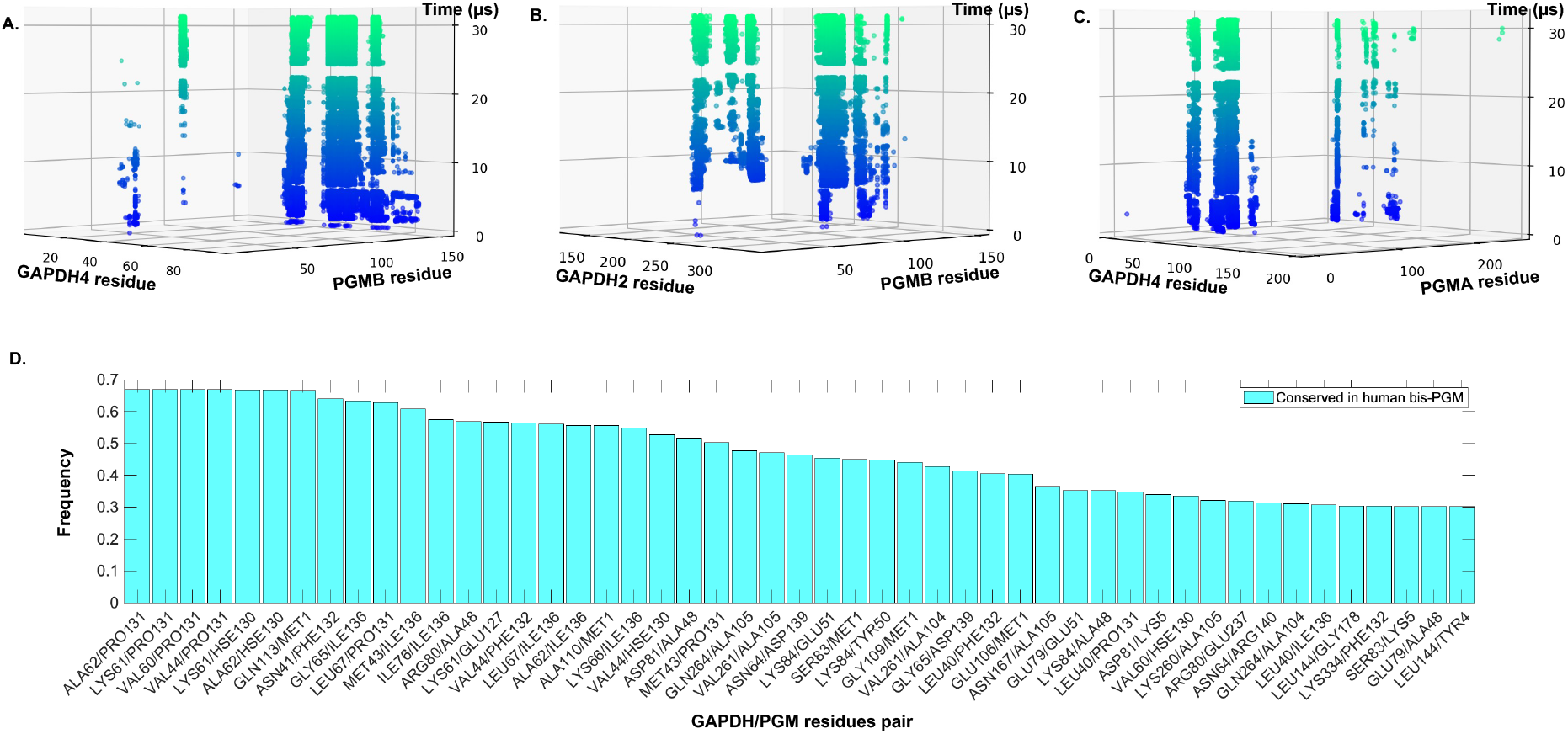
Residue specific contacts formed between human PGM and GAPDH in the mammalian cytoplasm simulation. Time traces for residue-residue contacts formed between (A) PGM subunit B and GAPDH subunit 4; (B) PGM subunit B and GAPDH subunit 2; and (C) PGM subunit A and GAPDH subunit 4. Each contact is shown as a circle, undergoing color change from blue to light green with increasing time. The distance cutoff for the residue-residue contacts in (A)-(C) was 0.45 nm. (D) Residue-residue contact frequency bar chart for the most frequent (≥30% of total time) interactions between PGM and GAPDH. PGM/GAPDH enzyme subunits pairs were included in the analyses if they had a total interacting area of ≥ 87.9 nm^2^, a cutoff picked to reflect the most significant interactions. The GAPDH/PGM residue pairs shown in the bar-chart involve PGM residues conserved across erythrocyte bis-PGM (with ∼53% sequence identity). See SI for all significant PGM/GAPDH subunits pairs, including non-conserved residues between PGM and bis-PGM, as discussed in the main text. A PGM residue is marked here as conserved if the corresponding bis-PGM residue either is same or similar. We grouped similar amino acids as following: GAVLI, FYW, CM, ST, KRH, DE, and NQ.

### Concerted disruption of the H-bond network when a quinary interaction is broken

Part of the PGK patch running from residues 130 to 143, comprised of Lys138, Ala139, Ser140, Glu142, and Asp143, is seen to form an intricate network of hydrogen bonds with GAPDH2’s Gln113, Tyr140, and Lys145 residues in Fig. 6 at Δt=0. The remaining three frames in Fig. 6 show the disruption of that hydrogen bond network, in 24 ns steps up to 22.9 μs (arrow in Fig. 4A). The breaking of the hydrogen bonds between GAPDH2/PGK in Fig. 6 is a concerted process: once the GAPDH2 Tyr140 interaction with the PGK sub-patch is disrupted at Δt=24ns, GAPDH2 Gln113 follows quickly at Δt=48 ns, and finally Lys145 at Δt=96 ns (not shown) completes the disruption. Concerted water penetration into the region between side chains facilitates the hydrogen bond network breaking. A case in point is the interaction between PGK Glu142 and GAPDH2 Lys145. We observe that patches form and dissolve contacts cooperatively, with a small network of water molecules moving in or out swiftly once the process of quinary contact formation or dissolution has started.

**Fig. 6.**
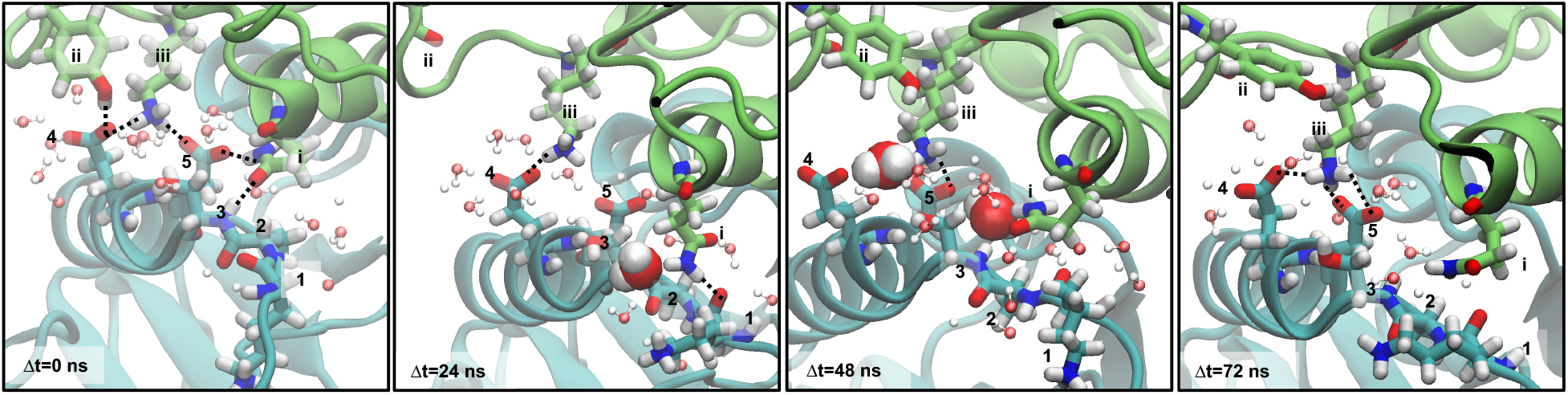
Hydrogen bond network being formed and broken between a patch of sidechains on the yeast PGK (N-terminal domain) and the human GAPDH (subunit 2) during simulation. Residues numbered 1-5 are on PGK: **1**. Lys138; **2**. Ala139; **3**. Ser140; **4**. Glu142; and **5**. Asp143. Residues numbered i-iii are on GADPH: **i**. Gln113; **ii**. Tyr140; and **iii**. Lys145. Structural representations as colored as follows: PGK backbone (cyan colored cartoon); GAPDH (lime colored cartoon); water molecule within 0.35 nm of both a numbered PGK residue and a numbered GAPDH residue (translucent CPK model); and licorice sticks (numbered residues with atoms shown as red oxygen, blue nitrogen, white hydrogen, cyan carbon if on PGK, and lime carbon if on GAPDH). Certain water molecules are further highlighted as the large space-filling VDW models when they either bridge or are displaced between a PGK/GAPDH residue pair from one frame to the next. A. The 1^st^ frame (on the outermost left) is at 22.8 μs of MD simulation and each subsequent frame is shown at a 24 ns time interval Δt later. 24 ns following the 4^th^ frame, the interaction between PGK and the GAPDH subunit has been completely broken, as can also be seen in the break between the dark and medium green circle columns in Fig. 4A.

### Hinge-bending motions in PGK

So far, we have focused on the intermolecular contacts between GAPDH, PGK, and PGM. PGK is itself a two-domain enzyme with N- and C-terminal domains connected by a hinge (cyan structure in Fig. 1). It has been surmised that opening of the hinge likely facilitates placement of the two substrates (ADP and 1,3-bisphosphoglycerate) on the two halves of the PGK active site, whereas closing of the hinge brings the two sites together to enable catalysis. The closed conformation allows phosphate to be channeled from the metabolic substrate on the N-terminal domain to the ADP bound to the C-terminal domain^38–40^. Coarse-grained simulation and experiments have shown that the hinge is more closed in crowded environments^7^. There is no high-resolution published structure yet of a closed conformation of yeast PGK.

We observe such intramolecular contact formation in our simulation. In Fig. 7, PGK shifts from an open state at ∼105° bending angle towards a compact structure with an approximately 90° bending angle. Simultaneous presence of a bound substrate on the N-terminal domain and a bound substrate on the C-terminal domain has been proposed to physiologically trigger the closed conformation^39,40^. In our simulations, the conformational change occurred without full active site binding. The substrate of GAPDH, glyceraldehyde-3-phosphate, was seen, however, to transiently bind PGK at a region around the N-terminal domain and the inter-domain from 1.58 μs to 1.97 μs. However, neither ADP nor ATP was seen binding to the C-terminal domain. In summary, PGK appears to close spontaneously, as proposed in ref. ^7^, although our simulation time is not sufficiently long to verify an open-closed equilibrium that favors the closed conformation for the ligand-free state of PGK.

**Fig. 7.**
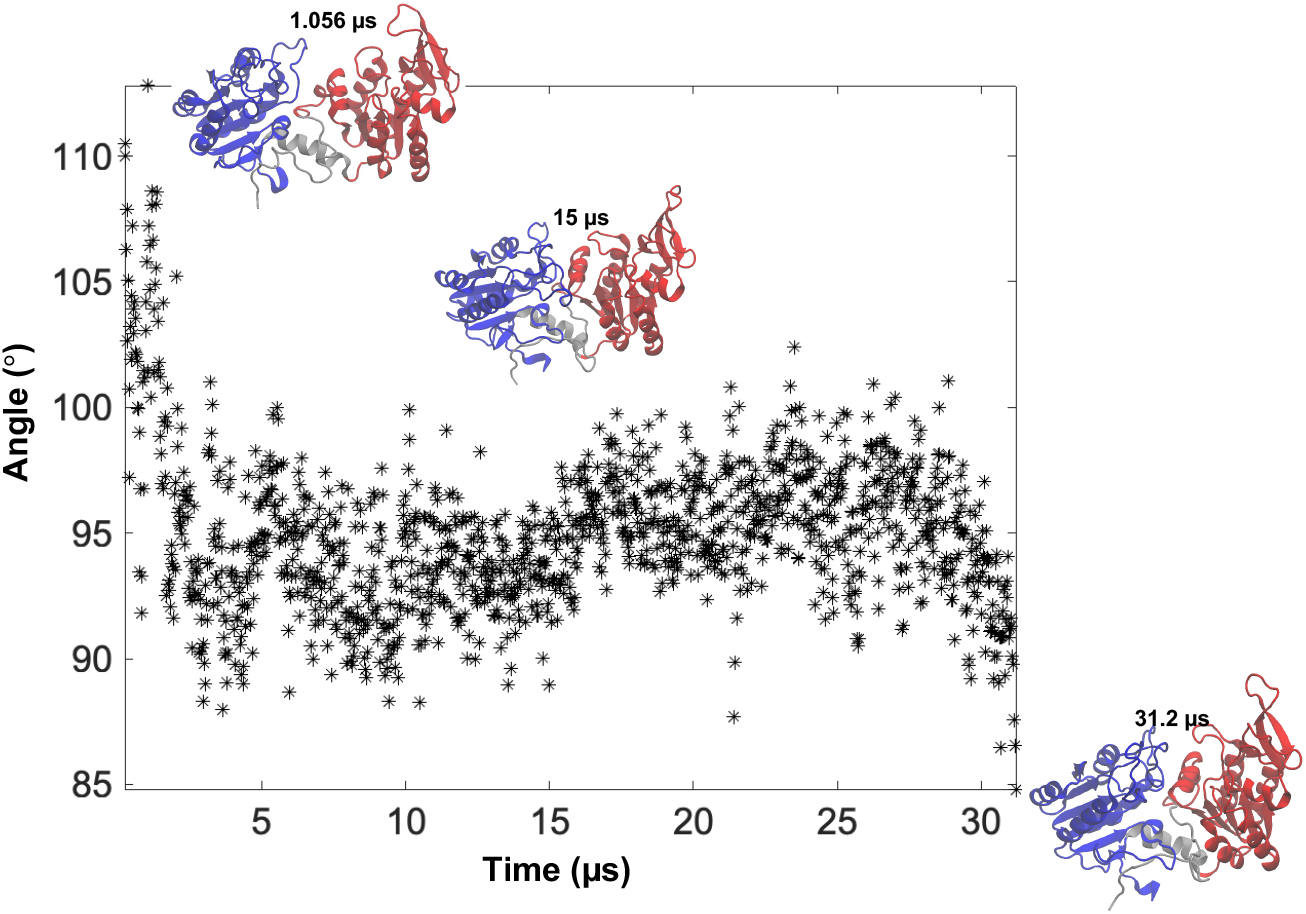
Hinge-bending motion in yeast PGK structure shifts the PGK conformation from “open” to “closed” during the simulation. The inter-domain angle was calculated as the angle between the center-of-mass of the N-terminal domain (residues 1-184), linker region connecting the N- and C-terminal domains (residues 185-199), and C-terminal domain (residues 200-393). Cartoon figures represent snapshots of PGK backbone at 3 different simulation time points across varying states of compactness with the interdomain angle at 1.056 μs being 112.8°, at 15 μs being 89°, and at 31.2 μs being 84.8°. PGK cartoons show N-terminal domain in blue, C-terminal domain in red, and inter-domain regions (residues185-199 and 394-415) in silver.

## Conclusion

In conclusion, we see that successive enzymes in a biochemical cycle form transient contacts in the cytoplasmic environment, lasting 1 to 5 μs as they break and re-form. These contacts occur as surface patches that form quinary structure, and contacts between these patches form and dissolve in a cooperative manner as several water molecules edge in or out between side chains. Not only do intermolecular contacts form, but the cytoplasm also promotes spontaneous formation of an intermolecular contact between the two domains of PGK, bringing the two halves of its active site in proximity, in our case without substrate binding. This model could form a scaffold for QM/MM calculations of enzymatic activity of PGK as ADP and 1,3-bisphosphoglycerate fall in place and are brought together.

## Methods

### Cytoplasm model construction and simulation

Model building and of the human cytoplasm model followed the general steps pursued for our previously published bacterial cytoplasm model^20^. The cytoplasm model energy was minimized for 6000 steps after solvation, pre-equilibrated according to previous methods^20^, and equilibrated briefly (∼7.3 ns) using NAMD 2.9 ^41^ without constraints before being transferred for extensive production runs on Anton 2 using the Charmm36m force field. We previously showed that this force-field produces less macromolecular sticking than Charmm22*^20^. The model was simulated for 31.2 μs using an NPT ensemble with a multigrator^42^ scheme at 1 atm pressure and 318 K. This temperature was selected to maximize protein diffusion while staying within realistic bounds for the cellular environment.

### Data analysis

A snapshot of the 31.2 μs trajectory was taken every 24 ns and the initial 240 ns of the trajectory was not considered for data analysis. Simulation trajectories were wrapped with VMD’s PBCTools plugin prior to data analysis to address periodic boundary conditions when calculating inter-residue contact distances. Data analysis of trajectories was done with both VMD^43^ and CPPTRAJ^44,45^ software packages.

## Acknowledgements

Anton 2 computer time was provided by the Pittsburgh Supercomputing Center (PSC) through Grant R01GM116961 from the National Institutes of Health. The Anton 2 machine at PSC was generously made available by D.E. Shaw Research. T.V.P. acknowledges support from the NIH grant R01-GM141298 and the Department of Chemistry, University of Illinois at Urbana-Champaign. P.S.R., M.R., and M.G. were supported by the NSF grant MCB 2205665.

## AUTHOR INFORMATION

The authors declare no competing financial interests.

## SI Figures and Tables

**Fig. S1.**
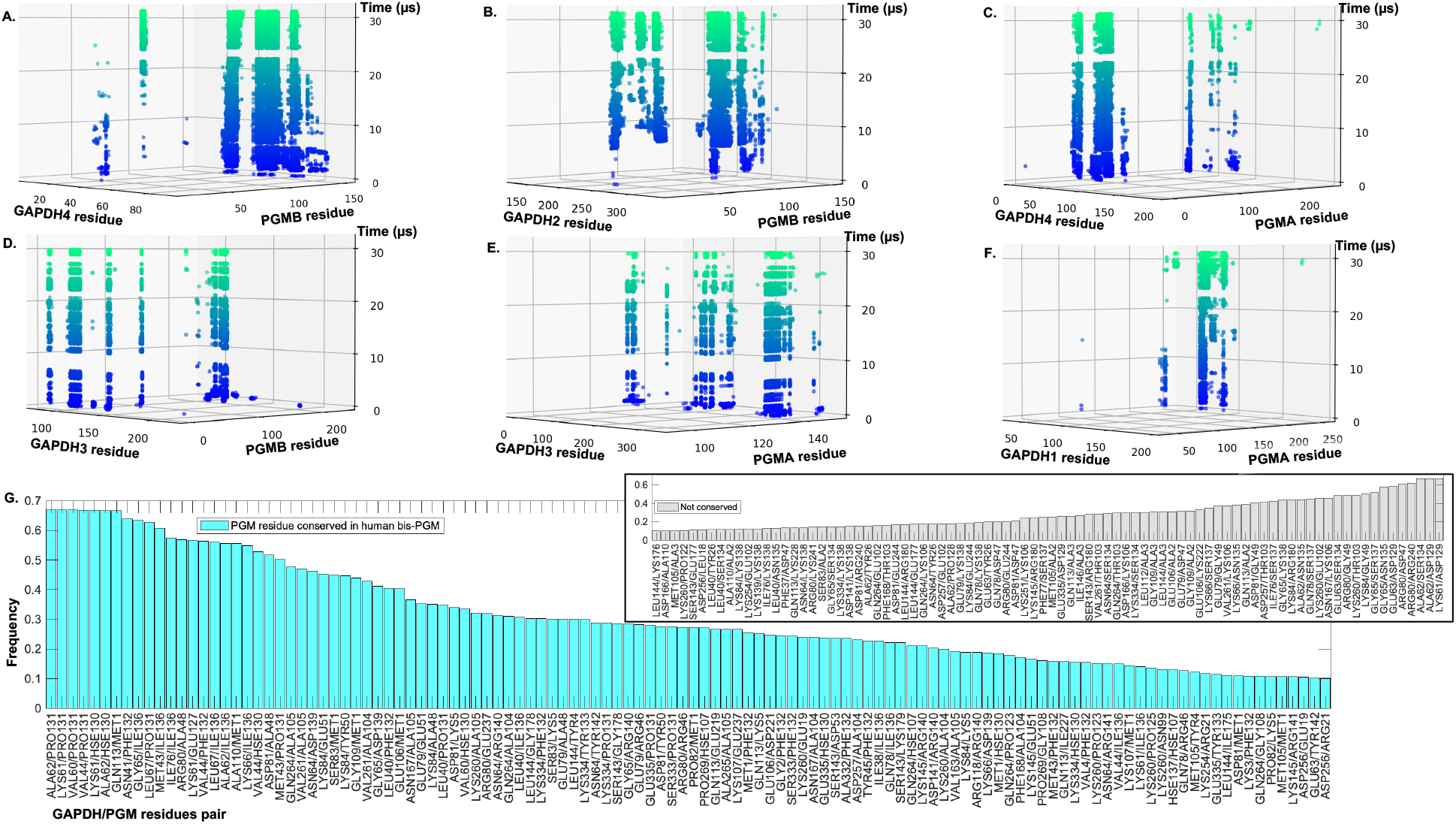
Full set of residue specific contacts formed between human PGM and GAPDH in the mammalian cytoplasm simulation. The distance cutoff for the residue-residue contacts in (A)-(F) was 0.45 nm. Residue-residue contact frequency bar charts is shown for the most frequent (≥10% of total time) interactions between PGM and GAPDH. PGM/GAPDH enzyme subunits pairs were included in the analyses in this figure if they had a total interacting area of ≥ 87.9 nm^2^, a cutoff picked to reflect the most significant interactions. The bar plots are colored cyan when there is enzyme residue conservation between PGM and its other human homolog, the erythrocytic bis-PGM (with 53% sequence identity); and white, when no PGM/bis-PGM residue conservation. A PGM residue is marked here as conserved if the corresponding bis-PGM residue either is same or similar. We grouped similar amino acids as following: GAVLI, FYW, CM, ST, KRH, DE, and NQ.

**Table S1.** Consensus abundances of metabolites in human cells, and number of each metabolite in the cytoplasm model chosen from the list by Monte Carlo sampling.

| Metabolite Name | Experimental Concentration (mM) |  | Consensus Concentration (mM) | No. of Copies Expected in Model | No. of Copies in Model | Concentration in Model (mM) |
| --- | --- | --- | --- | --- | --- | --- |
|  | *Chen et al.*<sup>+</sup> | *Diener et al.* |  |  |  |  |
| Glutamate | 24.628 | 39.138 | 31.883 | 42.2 | 41 | 31.000 |
| Taurine | 18.403 | 0.000 | 16.901 | 22.4 | 25 | 18.902 |
| Aspartate | 6.219 | 25.248 | 15.733 | 20.8 | 21 | 15.878 |
| Glutamine | 1.369 | 30.933 | 11.006 | 14.6 | 13 | 9.829 |
| Glycine | 6.725 | 13.500 | 10.112 | 13.4 | 13 | 9.829 |
| GSH | 5.551 | 10.865 | 8.208 | 10.9 | 6 | 4.537 |
| Spermine | - | 7.240 | 7.240 | 9.6 | 5 | 3.780 |
| Lactate | 1.874 | 11.310 | 5.461 | 7.2 | 5 | 3.780 |
| Asparagine | 0.145 | 7.290 | 3.717 | 4.9 | 2 | 1.512 |
| Malate | 0.679 | 5.273 | 3.651 | 4.8 | 10 | 7.561 |
| Alanine | 3.330 | 3.704 | 3.517 | 4.7 | 6 | 4.537 |
| Spermidine | - | 3.425 | 3.425 | 4.5 | 2 | 1.512 |
| Beta-Alanine | 6.481 | 0.329 | 3.405 | 4.5 | 4 | 3.024 |
| Citrate/Isocitrate | 1.329 | 4.904 | 3.117 | 4.1 | 7 | 5.293 |
| Hydroxyproline | - | 2.707 | 2.707 | 3.6 | 5 | 3.780 |
| ATP | 4.641 | 0.311 | 2.476 | 3.3 | 2 | 1.512 |
| Putrescine | - | 2.084 | 2.084 | 2.8 | 1 | 0.756 |
| Phosphocholine | 3.816 | 0.000 | 1.908 | 2.5 | 3 | 2.268 |
| UTP | 1.840 | 0.000 | 1.840 | 2.4 | 4 | 3.024 |
| Isoleucine | 1.401 | 3.255 | 1.709 | 2.3 | 1 | 0.756 |
| NAD | 0.489 | 2.712 | 1.601 | 2.1 | 2 | 1.512 |
| Serine | 0.471 | 4.031 | 1.588 | 2.1 | 2 | 1.512 |
| Leucine | 1.011 | 3.257 | 1.572 | 2.1 | 1 | 0.756 |
| Threonine | 1.005 | 2.189 | 1.278 | 1.7 | 2 | 1.512 |
| Arginine | 0.186 | 3.270 | 1.234 | 1.6 | 4 | 3.024 |
| Histidine | 1.122 | 1.073 | 1.098 | 1.5 | 1 | 0.756 |
| Valine | 1.348 | 1.397 | 1.096 | 1.5 | 1 | 0.756 |
| GSSG | 0.064 | 2.048 | 1.056 | 1.4 | 1 | 0.756 |
| UDP | 0.444 | 1.264 | 0.854 | 1.1 | 1 | 0.756 |
| Tyrosine | 0.678 | 1.497 | 0.838 | 1.1 | 1 | 0.756 |
| 3-Hydroxybutyric acid | - | 0.059 | 0.680 | 0.9 | 2 | 1.512 |
| G3P | 0.000 | 1.324 | 0.662 | 0.9 | 1 | 0.756 |
| Fumarate | 0.070 | 1.175 | 0.622 | 0.8 | 1 | 0.756 |
| GTP | 0.600 | 0.000 | 0.600 | 0.8 | 1 | 0.756 |
| ADP | 0.140 | 0.834 | 0.487 | 0.6 | 2 | 1.512 |
| AMP | 0.023 | 0.898 | 0.460 | 0.6 | 1 | 0.756 |
| FBP | 0.807 | 0.064 | 0.436 | 0.6 | 1 | 0.756 |
| Succinate | 0.042 | 0.774 | 0.408 | 0.5 | 1 | 0.756 |
| Pyruvate | 0.281 | 0.000 | 0.321 | 0.4 | 1 | 0.756 |
| R5P/RL5P | 0.114 | 0.374 | 0.244 | 0.3 | 1 | 0.756 |
| Pantothenic Acid | 0.235 | 0.000 | 0.235 | 0.3 | 1 | 0.756 |
| G6P | 0.146 | 0.322 | 0.234 | 0.3 | 1 | 0.756 |
| Citrulline | 0.009 | 0.634 | 0.217 | 0.3 | 1 | 0.756 |
| GMP | 0.111 | 0.079 | 0.095 | 0.1 | 1 | 0.756 |

\*The references to Chen *et al.* and Diener *et al.* in Table 1 are given in the main text.

## Notes

### Competing Interest Statement

The authors have declared no competing interest.

